# Dual roles of histone H3 lysine-4 in antagonizing Polycomb group function and promoting target gene expression

**DOI:** 10.1101/2024.06.25.600669

**Authors:** Cyril S. Anyetei-Anum, Mary P. Leatham-Jensen, Geoffrey C. Fox, B. Rutledge Smith, Krzysztof Krajewski, Brian D. Strahl, Jill M. Dowen, A. Gregory Matera, Robert J. Duronio, Daniel J. McKay

## Abstract

Tight control over cell identity gene expression is necessary for proper adult form and function. The opposing activities of Polycomb and trithorax complexes determine the ON/OFF state of targets like the Hox genes. Trithorax encodes a methyltransferase specific to histone H3 lysine-4 (H3K4). However, there is no direct evidence that H3K4 regulates Polycomb group target genes *in vivo*. Here, we demonstrate two key roles for replication-dependent histone H3.2K4 in target control. We find that H3.2K4 antagonizes Polycomb group catalytic activity and that it is required for proper target gene activation. We conclude that H3.2K4 directly regulates expression of Polycomb targets.

## Introduction

The characteristic size and shape of animal body parts is determined by the activity of master regulator genes during development. Either loss of function or ectopic expression of master regulators can result in identity changes, most recognizably demonstrated by the homeotic transformations caused by mutations of the *Drosophila* Hox genes, *Ultrabithorax* and *Antennapedia* in *Drosophila*. Due to their distinctive power to control form and identity, expression of master regulator genes is tightly controlled.

The Polycomb and trithorax groups of epigenetic regulators provide a crucial layer of control over master regulator gene expression, and the balance between their activities determines whether target genes are expressed. Polycomb group factors are required for heritable silencing of target genes (Kuroda et al. 2020). Multiple Polycomb group complexes exist, broadly categorized into two Polycomb Repressive Complexes, PRC1 and PRC2. Both PRC1 and PRC2 catalytically modify histones, and both can also recognize and bind to these post-translational modifications (PTMs). PRC2 tri-methylates histone H3 lysine-27 (H3K27me3), which is bound both by PRC1 (via Polycomb) and by PRC2 (via Esc) (Margueron et al. 2009). PRC1 ubiquitylates histone H2A lysine-119 (H2Aub1), which can be bound by factors associated with both PRC1 and PRC2 (Barbour et al. 2020). Binding of PRC2 to H3K27me3 leads to allosteric activation of the methyltransferase subunit, E(z). Through this allostery and other positive feedback mechanisms, PRC complexes spread across target genes to create repressive chromatin domains during initiation of Polycomb group target gene repression (Kuroda et al. 2020). These feedback loops also contribute to the maintenance of repressive chromatin domains following incorporation of unmodified nucleosomes during DNA replication, thus leading to stable propagation of transcriptionally silent states through rounds of cell proliferation.

Trithorax group factors counteract Polycomb group-mediated repression of target genes (Kingston and Tamkun 2014). First identified as suppressors of dominant Polycomb group mutant phenotypes, trithorax group loss of function results in loss of master regulator gene expression. Like Polycomb group complexes, many trithorax group proteins directly bind target genes. Although initial models proposed that trithorax group proteins function as transcriptional activators, subsequent genetic analyses indicated that trithorax group proteins may function as anti-repressors. Cells in which trithorax group and Polycomb group genes are both mutated still express targets such as the Hox genes (Klymenko and Müller 2004), suggesting that trithorax group proteins are not required for gene activation *per se*.

The mechanisms by which trithorax group proteins antagonize Polycomb group-mediated target gene repression remain unclear, in part because trithorax group proteins exhibit a wide range of biochemical activities. Most trithorax group genes encode chromatin-associated proteins with roles in gene regulation (Kingston and Tamkun 2014). These include the ATP-dependent nucleosome remodeler Kismet and multiple members of the SWI/SNF complex. Other trithorax group members encode histone methyltransferases, including the eponymous member, Trithorax, which methylates H3K4 (Tie et al. 2014), as well as Ash1, which methylates H3K36 (Yuan et al. 2011).

Several lines of evidence support a role for methylation of H3K4 and H3K36 in antagonizing Polycomb group function. *In vitro* biochemical studies have found that nucleosomes containing either H3K36me3 or H3K4me3 inhibit methylation of H3K27 by PRC2 (Schmitges et al. 2011; Voigt et al. 2012; Finogenova et al. 2020). Mass spectrometry results indicate that H3K36me3 or H3K4me3 are rarely found on the same tail as H3K27me3 (Voigt et al. 2012; Yuan et al. 2011). Consistent with these findings, methylation of H3K27 by PRC2 is only inhibited when H3K36me3 or H3K4me3 are present on the same tail (i.e. in *cis*) (Schmitges et al. 2011; Voigt et al. 2012; Yuan et al. 2011). Genetic evidence recently obtained in *Drosophila* now supports a direct role for H3K36 modification in inhibiting Polycomb group-mediated repression of Hox genes *in vivo* (Finogenova et al. 2020; Salzler et al. 2023). However, support for the role of H3K4 in Polycomb group target gene regulation remains equivocal (Dorafshan et al. 2019).

Chromatin profiling studies have determined that H3K4me1 is enriched at Polycomb Response Elements (PREs) (Rickels et al. 2016), the *cis*-elements that help recruit Polycomb group complexes to target genes in *Drosophila*, supporting the potential for H3K4me to directly antagonize Polycomb group function. However, despite global reduction of H3K4me1 levels, catalytically-deficient mutants of the H3K4 mono-methyltransferase Trr are viable and fertile without evidence of Polycomb group target gene dysregulation (Rickels et al. 2017). Catalytically-deficient mutants of the Trr orthologs *MLL3/4* likewise fail to exhibit Polycomb group target gene dysregulation in mouse embryonic stems cells or in developing embryos (Rickels et al. 2017; Xie et al. 2023; Dorighi et al. 2017). On the other hand, catalytic-deficient mutations of the H3K4 mono-methyltransferase Trx suppress Polycomb group mutant phenotypes in *Drosophila*, leaving open the possibility that selective recruitment of Trx inhibits Polycomb group function (Tie et al. 2014). It is unclear whether this antagonism is due to H3K4 or to non-histone substrates of Trx (Poreba et al. 2022). Consistent with the possibility of a non-histone target of Trx, H3K4 mutant cells were reported to exhibit no defects in the expression of genes downstream of major developmental signaling pathways in *Drosophila* (Hödl and Basler 2012), arguing against a direct role for H3K4 methylation in Polycomb group target gene regulation.

Here, we examine the role of H3K4 in control of Polycomb group target gene regulation in *Drosophila* development. Using a histone replacement system that changes replication-dependent H3.2K4 to H3.2K4R, we provide evidence that H3K4 methylation functions in two ways: to antagonizes PRC2 function while simultaneously contributing to the activation of Polycomb group targets, including the Hox genes.

## Results and Discussion

### H3.2K4 is required for completion of development

Study of epigenetic mechanisms often involves mutational analysis of histone-modifying enzymes that catalyze histone PTMs. However, these enzymes typically have non-histone substrates and non-catalytic functions (Morgan and Shilatifard 2023), obscuring the true effectors of epigenetic phenomena. An alternative strategy is to mutate the histone genes themselves. This approach is feasible in *Drosophila melanogaster* because the replication-dependent histone genes reside at a single genomic locus (Lifton et al. 1978), which can be deleted and complemented by transgenes bearing tandemly arrayed wild-type histone gene repeats (hereafter, *H3.2^WT^*) (Günesdogan et al. 2010; McKay et al. 2015). Mutagenesis of this histone replacement transgene eliminates the PTM site, thereby permitting direct interrogation of individual histone residue functions.

Prior examination of histone replacement genotypes in *Drosophila* reported the absence of transcriptional defects in H3K4 mutant cells (Hödl and Basler 2012). However, the viability of these mutants was not reported. To examine the developmental consequences of H3K4 mutation, we genetically replaced the endogenous replication-dependent histone H3.2 with an arginine substitution mutant (*H3.2^K4R^*) (**Figure 1A**, S1A). In comparison to *H3.2^WT^*controls, *H3.2^K4R^* replacement flies do not survive to adulthood (**Figure 1B**, Table S1). Examination of the lethal phase indicated that significantly fewer *H3.2^K4R^*mutant embryos hatch into larvae relative to *H3.2^WT^* controls (5% vs 90%) (**Figure 1C**, Table S1). Furthermore, the few *H3.2^K4R^* embryos that hatch into larvae arrest in early larval development (**Figure 1D**). Given the critical role of the H3K4 methyltransferase, Trx, in maintaining Hox gene expression and body segment identity (Breen and Harte 1993), we examined *H3.2^K4R^* embryos for morphological defects. Surprisingly, *H3.2^K4R^*mutant embryos show neither detectable changes in Ubx immunofluorescence signal nor defects in cuticle patterning (**Figures 1E**, S1B).

**Figure 1.**
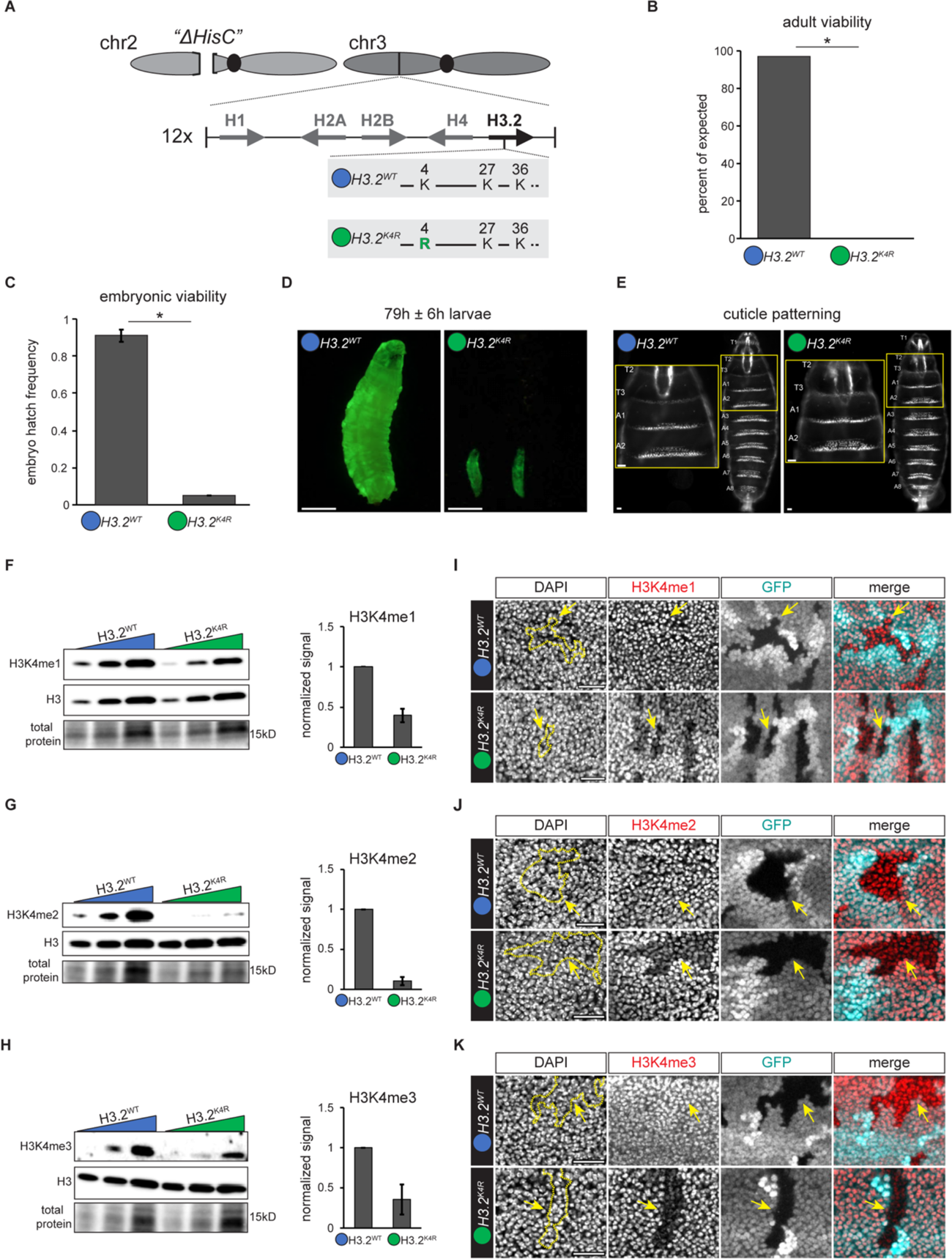
H3.2 K4 is required for completion of development. (A) Schematic of the histone gene replacement chromosomes. *ΔHisC* represents deletion of the endogenous replication-dependent histone genes. The *H3.2^WT^* (blue circle) and *H3.2^K4R^* (green circle) transgenes are shown. (B) Barplot of adult viability for the indicated genotypes plotted as percent of expected frequencies based on Mendelian ratios. Asterisk indicates p < 0.05 (chi-squared test). (C) Barplot of embryonic viability for the indicated genotypes. Histone replacement transgenes inherited maternally. Error bars indicate ±SEM. Asterisk indicates p < 0.05 (student’s t-test). (D) Comparison of *H3.2^WT^* and *H3.2^K4R^* larvae. Scale bar indicates 1mm. Histone replacement transgene inherited paternally. (E) Image of *H3.2^WT^* and *H3.2^K4R^* cuticles. Inset highlighted with yellow box. Scale bar indicates 20µM. (F-H) Western blots of H3K4me1 (F), H3K4me2 (G), and H3K4me3 (H) from 16-18h old *H3.2^WT^* and *H3.2^K4R^* embryos. Barplots depict average fold change in H3K4me signal normalized to total protein and plotted relative to *H3.2^WT^*. Error bars represent ±SEM. At least three biological replicates were performed for western blots. (I-K) Confocal images of *H3.2^WT^* and *H3.2^K4R^* mitotic clones in wing imaginal discs stained for DAPI (gray), H3K4me1 (I), H3K4me2 (J), or H3K4me3 (K) (red), and GFP (cyan). Merge includes H3K4me and GFP signals. Dashed line outlines clone highlighted with yellow arrows. Scale bar indicates 20 µm.

To assess H3K4 methylation levels in late-stage embryos, we performed western blotting and observed a 2.5-fold decrease in H3K4me1, a 9.6-fold decrease in H3K4me2, and a 2.8-fold decrease in H3K4me3, in *H3.2^K4R^* embryos relative to *H3.2^WT^* control embryos (**Figure 1F–H**). Residual H3K4me1-me3 western blot signals may be due to expression of the replication-independent *H3.3* genes, which remain intact in these genotypes. *H3.2^K4R^* mutant embryos also have remaining wild-type maternal H3.2 histones that package the genome during nuclear cycles preceding zygotic histone gene expression (Marzluff et al. 2008). This maternal H3.2 may also contribute to the lack of morphological defects observed in *H3.2^K4R^* embryos. Consistent with this interpretation, the hatching frequency of *H3.2^K4R^* embryos that maternally inherit the mutant transgene, which possess a mixture of wild-type and H3.2K4R mutant histones, is lower than those that paternally inherit the mutant transgene, which express the mutant transgene only after zygotic genome activation (5% vs 20%, respectively) (Figures S1C-H). We conclude that H3.2K4 is required for completion of development.

### H3.2K4 is not required for enhancer or gene activation

Due to the lack of obvious morphological defects in *H3.2^K4R^* mutant embryos, we turned to genetic mosaic analysis in proliferating larval tissues to further examine the role of H3.2K4 in developmental gene regulation. In these experiments, all copies of the wild-type, replication-dependent histone genes are deleted via mitotic recombination (*ΔHisC* clones), leaving only transgenic histone gene expression. Similar to embryonic western blot results (Fig. 1F-H), we found that H3K4 methylation levels are reduced in *ΔHisC* clones expressing *H3.2^K4R^*mutant histones (hereafter, *H3.2^K4R^* clones), whereas H3K4me1-me3 levels were unchanged in control *H3.2^WT^* clones (**Figure 1I–K**). Given that H3K4 methylation is closely correlated with gene activity, we next examined the impact of the *H3.2^K4R^* mutation on gene expression and enhancer activation. Expression of the broadly expressed cell membrane protein, Fasciclin III, and the nuclear protein, Osa, are both unaffected in *H3.2^K4R^* clones (**Figure S2A, S2B**), consistent with prior findings that gene expression can still occur normally in H3K4 mutant cells (Hödl and Basler 2012).

We further explored the impact of *H3.2^K4R^* on activation of genes and enhancers that are transcriptionally inactive at the time of clone induction (mid-larval stages), reasoning that replication-dependent accumulation of H3.2K4R histones in target gene chromatin prior to transcription may interfere with gene activation. We examined an enhancer from the *nubbin* gene that is inactive in larval tissues, but which turns on in pupal wings (Uyehara et al. 2017). We also examined the ecdysone-induced transcription factor *E93*, which is likewise off in larval wing imaginal discs, but which turns on in pupal stages (Guo et al. 2016). The *E93* gene and the *nubbin* enhancer both exhibit low chromatin accessibility and low levels of H3.3 enrichment in wing imaginal discs **(Figure S2C, S2E)**, consistent with their transcriptional inactivity and lack of replication-independent histone incorporation at the time of *ΔHisC* clone induction (Salzler et al. 2023; Uyehara et al. 2017). We found that *H3.2^K4R^* clone induction in early wings does not impact activation of the *nubbin* enhancer or *E93* **(Figures S2D, S2F)**. We conclude that initiation of gene expression and activation of enhancers can occur normally in *H3.2^K4R^*cells.

### *H3.2^K4R^* replacement cells show decreased H3K27me2/3

Prior *in vitro* studies indicate that H3K4me3 inhibits methyltransferase activity of PRC2 (Kasinath et al. 2021; Schmitges et al. 2011). One potential consequence of *H3.2^K4R^* mutation may therefore be a loss of PRC2 inhibition, resulting in an increase in H3K27me3 levels. To examine the impact of *H3.2^K4R^* on H3K27 methylation *in vivo*, we performed immunofluorescence on *ΔHisC* clones in wing imaginal discs. Contrary to expectations, we found that *H3.2^K4R^* clones exhibit reduced rather than increased H3K27me3 (**Figure 2B**). A similar reduction in H3K27me2 was observed (**Figure 2A**). By contrast, H3K27 methylation levels were unchanged in control *H3.2^WT^* clones (**Figure 2A, 2B)**. Consistent with these findings, *H3.2^K4R^*embryos exhibit 4.6-fold reduced H3K27me3 levels relative to control *H3.2^WT^*embryos by western blot (**Figure 2C**). We observed no change in H2AK118ub1 levels in *H3.2^K4R^* clones **(Figure S2G)**, suggesting *H3.2^K4R^* selectively impacts PRC2 activity without impacting PRC1 catalytic activity.

**Figure 2.**
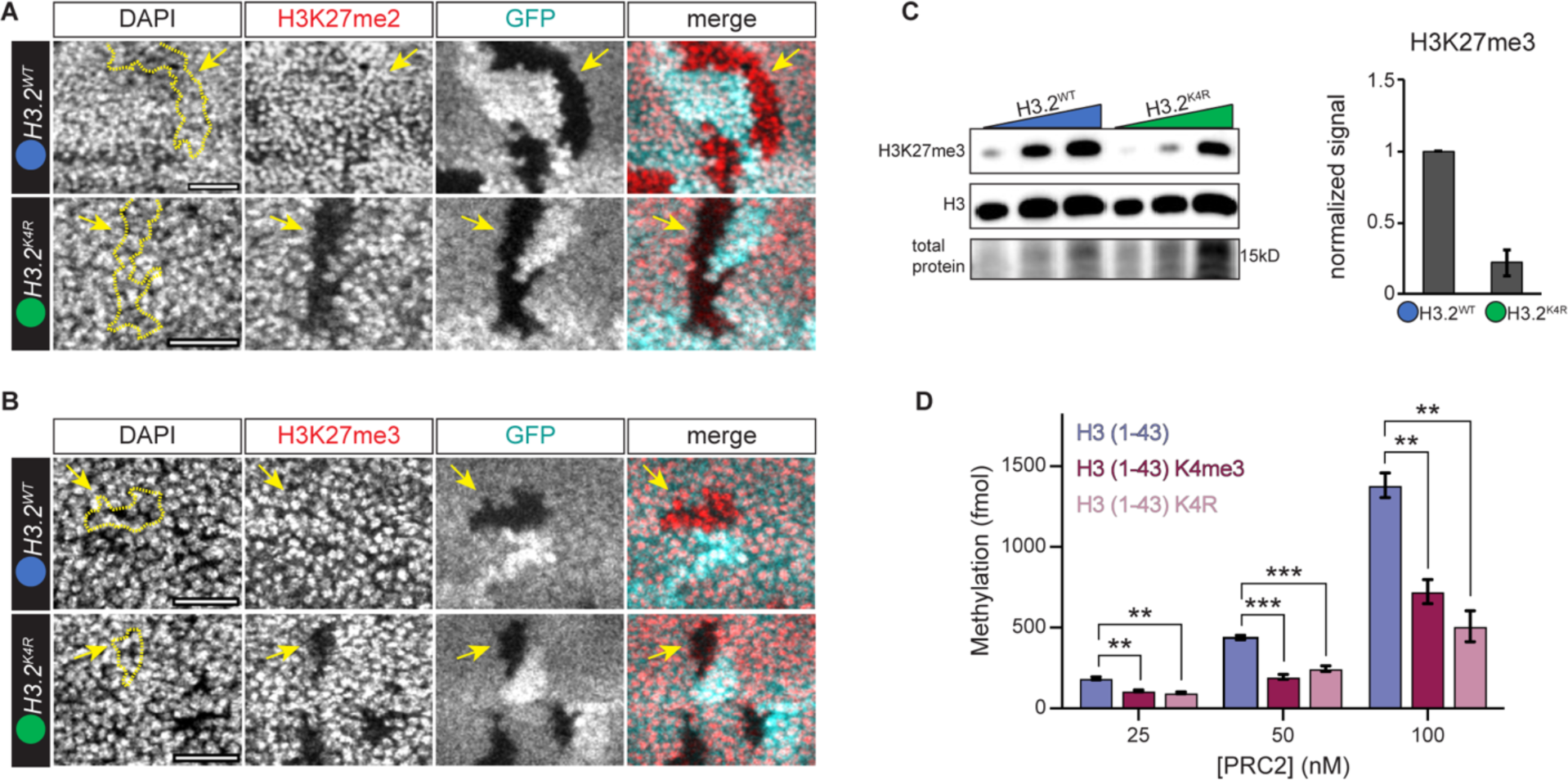
*H3.2^K4R^* replacement cells show decreased H3K27me2/3. (A-B) Confocal images of *H3.2^WT^* and *H3.2^K4R^* mitotic clones in wing imaginal discs stained for DAPI (gray), H3K27me2 (A), or H3K27me3 (B) (red), and GFP (cyan). Merge includes H3K27me and GFP signals. Dashed line outlines clone highlighted with yellow arrows. Scale bar indicates 20 µm. (C) Western blot of H3K27me3 from 16-18h old *H3.2^WT^* and *H3.2^K4R^* embryos. Barplot depicts average fold change in H3K27me3 signal normalized to total protein and plotted relative to *H3.2^WT^*. Error bars represent ±SEM. Three biological replicates were performed. (D) Barplot of average *in vitro* methylation signal for three concentrations of PRC2 on unmodified H3, H3K4me3, and H3K4R peptides (amino acids 1-43). Methylation activity is measured in femtomoles (fmol). Error bars indicate ±SEM. ** indicates p < 0.005, *** indicates p < 0.001 (student’s t-test).

A similar outcome was recently reported for H3.2K36, which like H3.2K4, inhibits PRC2 catalytic activity when methylated. These studies found that embryos expressing H3.2K36R mutant histones exhibit reduced H3K27me3 levels and de-repression of Polycomb group target genes *Ubx* and *Abd-B* (Finogenova et al. 2020; Salzler et al. 2023; Schmitges et al. 2011). Cryo-EM analyses indicated that mutation of H3.2K36 may disrupt the fit of H3.2K27 in the catalytic center of PRC2 in a manner similar to H3.2K36me3, providing an explanation for reduced levels of H3K27 methylation in *H3.2^K36R^* mutants (Finogenova et al. 2020), and raising the possibility that *H3.2^K4R^* mutants impact PRC2 activity in an analogous manner.

To test whether H3.2K4R also inhibits PRC2 catalytic activity, we performed *in vitro* methyltransferase assays on histone H3 peptides encoding the first 44 amino acids of the N-terminal tail. In agreement with prior studies, the high levels of PRC2-mediated methylation observed for unmodified H3 peptides is reduced an average of 2-fold across three concentrations of PRC2 when H3K4me3 peptides were used as substrate (**Figure 2D, S2H**). Intriguingly, H3.2K4R peptides yielded a similar 2.2-fold decrease in PRC2-mediated methylation (**Figures 2D, S2H).** This finding suggests that similar to H3.2K36R, H3.2K4R also disrupts the ability of the H3 tail to be read by PRC2 and hence, is acting similar to H3K4me3 by antagonizing PRC2 activity. Consistent with this interpretation, crystal structures of the histone H3.2 tail with the PRC2 complex member Caf1-55 indicate tight coordination of the H3.2K4 residue and predict steric interference by the addition of methyl groups to H3.2K4 (Schmitges et al. 2011). It is possible that the arginine side chain in H3.2K4R histones causes similar steric interference, resulting in reduced catalytic activity of PRC2. H3K4me3 has also been proposed to interact with the allosteric site on the PRC2 subunit EED (Cookis et al. 2024). Therefore, it is also possible that H3.2K4R similarly disrupts allosteric activation of PRC2. Regardless of the mechanism, our findings provide *in vivo* evidence that H3.2K4 counteracts PRC2-mediated H3.2K27 methylation.

### Repression of Polycomb group target genes is not impaired in *H3.2^K4R^* cells despite low H3K27me3 levels

Epigenetic silencing of Polycomb group target genes depends on H3K27 methylation (McKay et al. 2015; Pengelly et al. 2013). The reduction of H3K27 methylation observed in *H3.2^K4R^*cells prompted us to examine whether Polycomb group target genes become de-repressed in these cells. We focused on the Hox genes *Ubx* and *Abd-B*, which are both repressed by Polycomb group complexes in wing imaginal discs (**Figures 3A, S3)** (Beuchle et al. 2001). As expected, we observed no changes to H3K27me3 levels in *H3.2^WT^* clones, and both *Ubx* and *Abd-B* remain silenced (**Figure 3A-D**). In strong contrast, we observed a decrease in H3K27me3 to 26% of wild-type levels and robust de-repression of both *Ubx* and *Abd-B* in *H3.2^K27R^* wing imaginal disc clones (**Figures 3A-D**). We next examined *H3.2^K36R^* clones and observed a reduction in H3K27me3 to 60% of wild-type levels. Despite the reduction in H3K27me3 levels in *H3.2^K36R^* clones, we observed no de-repression of *Ubx* (**Figures 3A-D**). Examination of *H3.2^K4R^* clones revealed a decrease in H3K27me3 to 36% of wild-type levels. However, despite the strong decrease in H3K27me3 levels, we observed no de-repression of *Ubx* or *Abd-B* (**Figures 3A-D, S3)**. The lack of *Ubx* and *Abd-B* de-repression in *H3.2^K4R^* clones suggests that H3.2K4 methylation may play a critical role in Polycomb group target gene activation.

**Figure 3.**
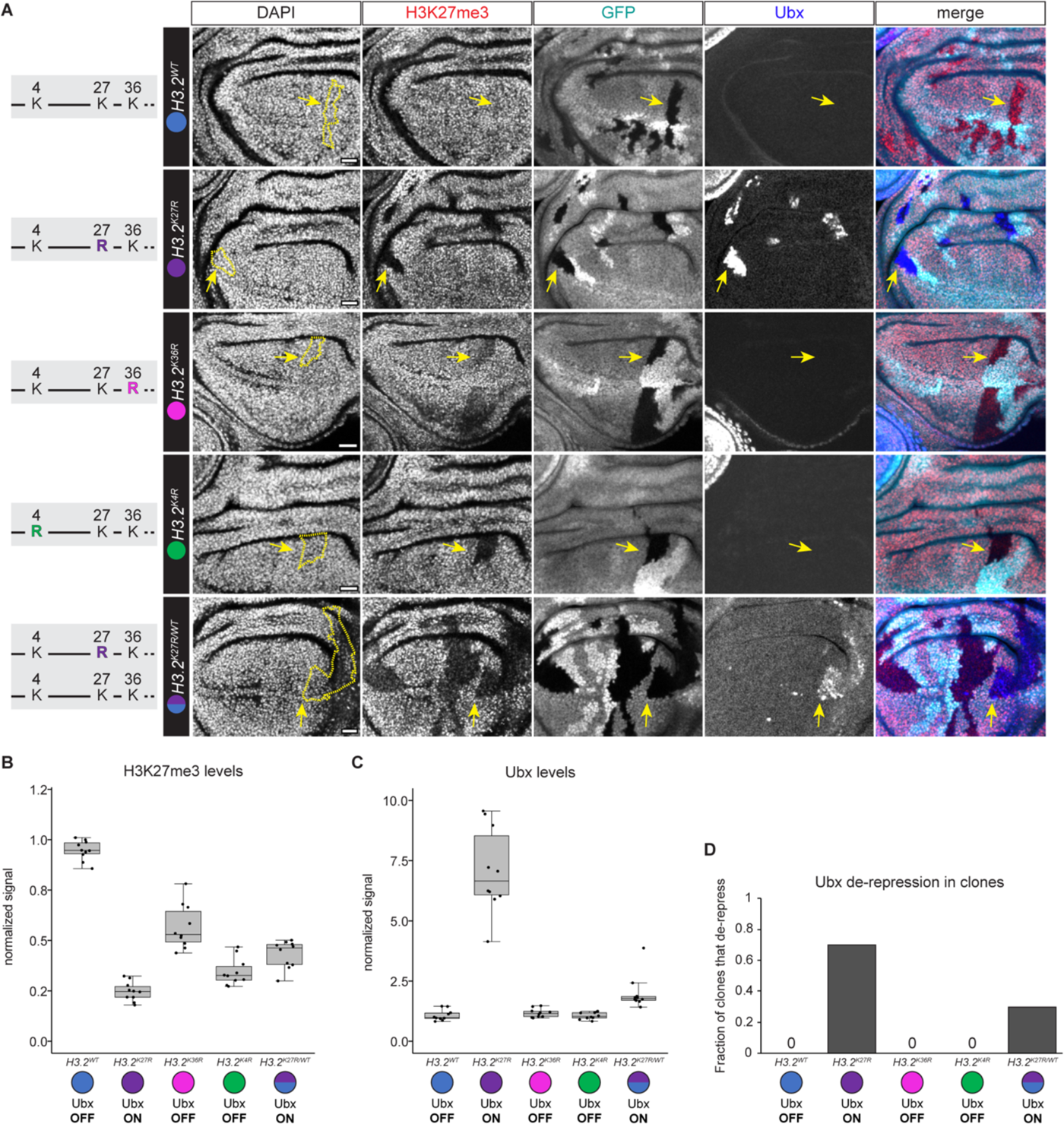
Repression of Polycomb group target genes is not impaired in *H3.2^K4R^* cells despite low H3K27me3 levels. (A) Confocal images of *H3.2^WT^, H3.2^K27R^, H3.2^K36R^, H3.2^K4R^, and H3.2^K27R/WT^* mitotic clones in wing imaginal discs stained for DAPI (gray), H3K27me3 (red), GFP (cyan), and Ubx (blue). Merge includes H3K27me3, GFP, and Ubx signals. Dashed line outlines clone highlighted with yellow arrows. Scale bar indicates 20 µm. (B, C) Boxplots of H3K27me3 (B) and Ubx (C) signal for the indicated genotypes plotted as the ratio of signal inside the clone normalized to signal outside the clone, plotted relative to *H3.2^WT^*. The box represents the interquartile range (IQR). The horizontal line indicates the median. Whiskers extend to the smallest and largest values within 1.5 times the IQR. Pairwise comparisons between all genotypes are statistically significant in (B). Pairwise comparisons between *H3.2^K27R^* or *H3.2^K4R,K27R^* and all other genotypes are statistically significant in (C), p < 0.05 (student’s t-test) (Table S2). (D) Barplot of the fraction of clones that de-repress Ubx for the indicated genotypes.

Because the ability of Polycomb group complexes to silence target genes closely depends on H3K27me3 levels (Coleman and Struhl 2017), we sought to determine if the H3K27me3 signal observed in *H3.2^K4R^* cells remained above a threshold sufficient for repression. To identify the de-repression threshold, we genetically defined the dose of wild-type and H3.2K27R mutant histones by creating a genotype that carried one *H3.2^WT^* transgene and one *H3.2^K27R^* mutant transgene (hereafter, *H3.2^K27R/WT^*) (**Figure 3A**). In this genotype, *ΔHisC* clones are expected to express a 1:1 ratio of wild-type and H3.2K27R histones. Consistent with this expectation, immunofluorescence indicated that H3K27me3 levels in *H3.2^K27R/WT^* clones are reduced to 46% of wild-type levels. More importantly, this reduction in H3K27me3 is accompanied by de-repression of *Ubx* (**Figures 3B, 3C)**. Only a subset of *H3.2^K27R/WT^* clones (30%) exhibit *Ubx* de-repression, and these are preferentially located in the posterior wing pouch region (**Figures 3A, 3D)**, consistent with prior reports of spatial influence over *Ubx* de-repression in wing imaginal discs (Beuchle et al. 2001). When de-repressed, the levels of *Ubx* expression are lower in *H3.2^K27R/WT^*relative to *H3.2^K27R^* clones, indicating that *Ubx* is only partially de-repressed (**Figure 3C**). Consistent with partial disruption of Polycomb group-mediated Hox gene repression, no *Abd-B* de-repression was observed in *H3.2^K27R/WT^* clones (Beuchle et al. 2001). Most importantly, H3K27me3 levels in *H3.2^K4R^* cells are significantly lower than in *H3.2^K27R/WT^* cells (**Figure 3B**), and despite these lower levels, *Ubx* is not de-repressed in *H3.2^K4R^*cells, unlike in *H3.2^K27R/WT^* cells (**Figures 3C, 3D)**. Collectively, these findings indicate that a reduction in H3K27me3 levels to ∼50% of wild-type levels is below the threshold amount needed for repression of *Ubx*. Thus, despite being below this threshold, *Ubx* is not de-repressed in *H3.2^K4R^* cells. These data argue that while H3.2K4R antagonizes PRC2 catalytic activity, it is also disrupting the normal function of this residue to promote gene expression.

### H3.2K4 promotes proper activation of Polycomb group target genes

One explanation for the lack of *Ubx* or *Abd-B* de-repression in *H3.2^K4R^* cells despite levels of H3K27me3 below that needed for repression is that H3.2K4 contributes to Polycomb group target gene activation. To investigate this possibility further, we generated a histone replacement transgene in which all H3.2 genes are doubly mutated at lysine-4 and lysine-27 (*H3.2^K4R,K27R^*). We reasoned that H3.2K27 mutation causes strong Polycomb group target gene de-repression and that comparison between *H3.2^K27R^*single mutant clones and *H3.2^K4R,K27R^* double mutant clones could reveal a role for H3.2K4 in Polycomb group target gene activation.

We observed that de-repression of some Polycomb group target genes is indeed compromised in *H3.2^K4R,K27R^* double mutant cells (**Figures 4A-D**). The levels of *Abd-B* de-repression in *H3.2^K4R,K27R^* double mutant clones are approximately 2-fold lower than the levels of *Abd-B* de-repression in *H3.2^K27R^* single mutant clones (**Figures 4A, 4B, and Table S2**). Likewise, de-repression levels of *abd-A*, another Polycomb group target gene in the Bithorax complex, are approximately 3-fold lower in *H3.2^K4R,K27R^* double mutant cells relative to *H3.2^K27R^* single mutant cells (**Figures 4C, 4D**). Interestingly, we did not observe a decrease in *Ubx* de-repression in *H3.2^K4R,K27R^* double mutant cells relative to *H3.2^K27R^* single mutant cells (**Figure 4E, 4F)**. On the contrary, *Ubx* levels significantly increased in *H3.2^K4R,K27R^*double mutant cells (**Figure 4F**). It is possible that the increase in *Ubx* levels in *H3.2^K4R,K27R^* double mutant cells is due to the observed decrease in *abd-A* and *Abd-B* expression. Posteriorly expressed Hox factors are known to repress expression of those that are more anteriorly expressed, and both *abd-A* and *Abd-B* repress *Ubx* (González-Reyes and Morata 1990).

**Figure 4.**
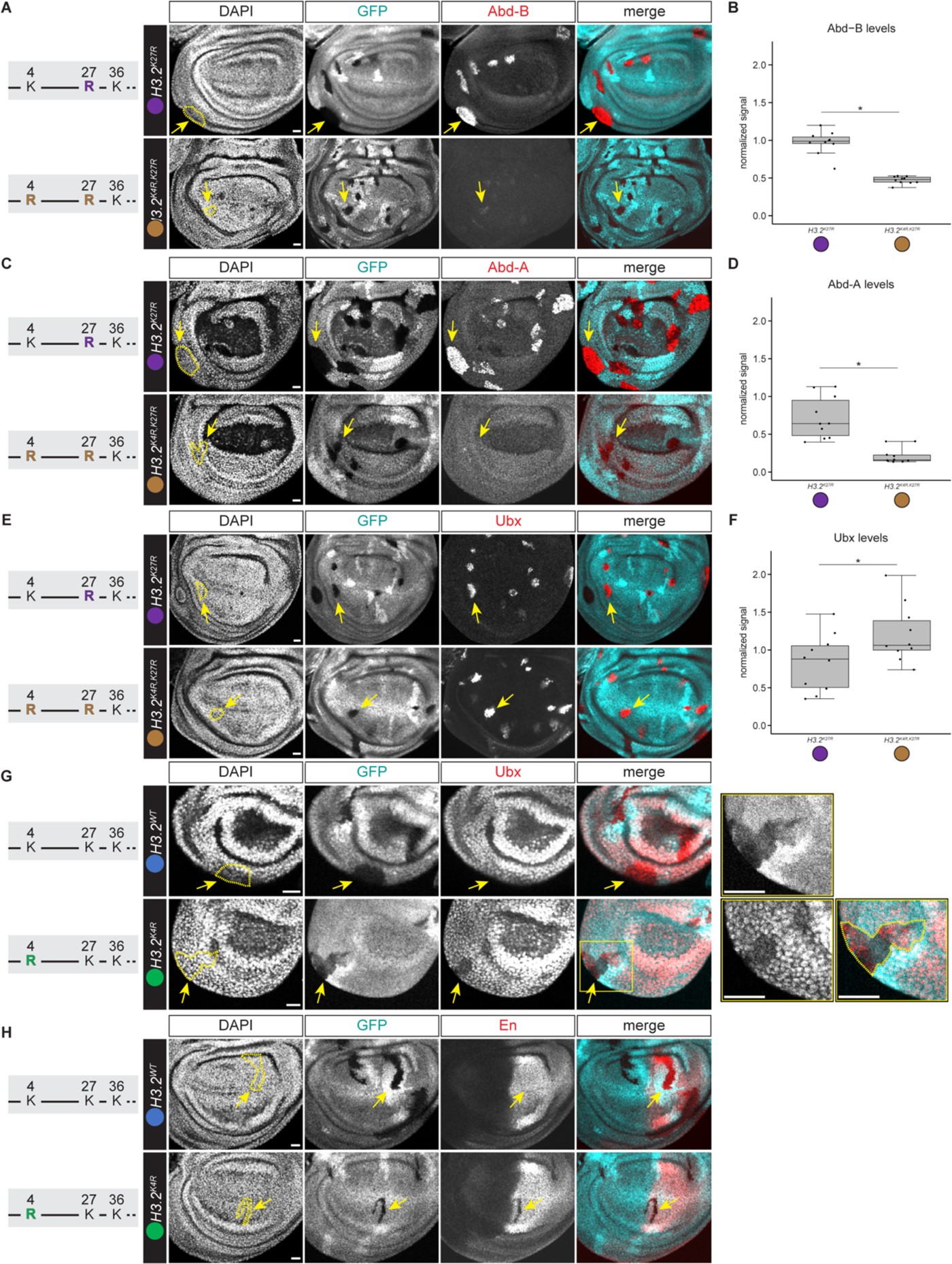
H3.2K4 promotes activation of Polycomb group target genes. (A, C, E, G, H) Confocal images of *H3.2^K27R^, H3.2^K4R,K27R^, H3.2^WT^, and H3.2^K4R^* mitotic clones in wing imaginal discs stained for DAPI (gray), GFP (cyan), Abd-B (A), Abd-A (C), Ubx (E), and En (H) (red). A haltere imaginal disc is shown in panel G with inset highlighted with yellow box. For all panels, scale bar indicates 20µM. Merge includes cyan and red channels. Dashed line outlines clone highlighted with yellow arrows. (B, D, F) Boxplots of Abd-B (B), Abd-A (D), and Ubx (F) signal for *H3.2^K27R^or H3.2^K4R,K27R^* plotted as the ratio of signal inside the clone normalized to signal outside the clone, plotted relative to *H3.2^K27R^*. The box represents the IQR. The horizontal line indicates the median. Whiskers extend to the smallest and largest values within 1.5 times the IQR. Asterisk indicates p < 0.05 (student’s t-test).

The findings presented above indicate that H3.2K4 contributes to Hox gene de-repression when Polycomb group function is disrupted by decreased H3.2K27me3. To ask whether H3.2K4 contributes to Polycomb group target gene activation during normal development, we examined expression of *Ubx* in *H3.2^K4R^*clones located in haltere and T3 leg imaginal discs, two tissues in which *Ubx* is normally expressed. We observed reduced *Ubx* expression in 32% of *H3.2^K4R^* clones (n=25 clones), whereas *Ubx* levels were unchanged in the remaining clones (**Figure 4G**). Similarly, expression of another Polycomb group target gene, *Engrailed* (*En*), was reduced in 50% of *H3.2^K4R^* clones (n=10 clones) (**Figure 4H**). Collectively, these observations indicate that H3.2K4 is required for proper maintenance of Polycomb group target gene expression during normal development.

In rare instances, we observed *H3.2^K4R^* clones in which *Ubx* expression appeared variegated (**Figure 4G**), with a subset of cells inside the clone exhibiting Ubx immunofluorescence signal and other cells inside the clone in which Ubx signal was absent. Because cells within a clone derive from a common progenitor cell that expressed *Ubx* at the time of clone induction, the appearance of variegated expression suggests that *Ubx* was silenced in a subset of the clone’s cells and subsequently maintained in the off state as these cells proliferated.

H3K4 methylation is one of the most broadly conserved and heavily studied histone PTMs, and its enrichment is closely correlated with active chromatin states. However, evidence supporting a causal role for H3K4 in gene regulation has remained elusive (Howe et al. 2017). Here, we provide strong support for two key functions of H3.2K4 in regulation of Polycomb group target gene expression. First, the findings demonstrate an *in vivo* role for H3.2K4 methylation in antagonizing PRC2 function, supporting prior *in vitro* evidence that PRC2 catalytic activity is inhibited by H3.2K4me3. We propose that the arginine substitution employed here functions analogously to H3.2K4me3 in antagonizing PRC2 function. A corollary to this interpretation is that unmodified H3.2K4 is important for enabling PRC2 catalytic activity *in vivo*. Future genomic profiling studies are needed to determine whether H3.2K4 is required to constrain the spread of Polycomb group chromatin domains, as has been proposed for H3K36 methylation (Yuan et al. 2011; Finogenova et al. 2020; Salzler et al. 2023). Second, reduced expression of Polycomb group target genes in *H3.2^K4R^*cells demonstrates a role for H3.2K4 methylation in promoting Polycomb group target gene expression. In contrast to PRC2 antagonism, we propose that the arginine substitution employed here does not function analogously to methylated H3.2K4 for gene activation. Instead, it is likely that methyl-H3.2K4 reader proteins do not bind H3K4R histones, disrupting their role in Polycomb target gene activation. Together, these findings place H3.2K4 as a focal point that balances two key epigenetic pathways controlling cellular memory.

Our findings reveal defective but not complete loss of Polycomb target gene activation in *H3.2^K4R^* cells, raising the question whether H3.2K4 methylation is absolutely required for their activation, or if it is one of multiple inputs. One important consideration of our mosaic analysis is the potentially confounding presence of wild-type parental histones inherited by the daughter cells following mitotic recombination. At the time of *ΔHisC* clone induction, the great majority of histones in chromatin are wild type, and these are inherited by daughter cells during DNA replication. As a result, the abundance of parentally contributed wild-type H3.2K4 may suffice to at least partially activate Polycomb target genes once H3.2K27me3 drops below the repression threshold. A second consideration is the presence of the replication-independent histone H3.3, which is enriched for H3K4 methylation (McKittrick et al. 2004), and which is still expressed in our experiments. Both these factors may explain why Polycomb target genes can still be activated in *H3.2^K4R^* cells despite H3K27me3 levels that are below the threshold needed for repression. The dependence on H3.2K4 for activation may also vary between Polycomb target genes. Indeed, *Ubx* de-repression occurs more readily than de-repression of *abd-A* or *Abd-B* in *H3.2^K4R,K27R^* cells, suggesting that *Ubx* is less dependent on H3.2K4 for activation. Additional studies, including development of new experimental approaches, are needed to resolve these questions.

The mechanisms by which H3.2K4 contributes to Polycomb group gene expression also remain to be determined. Multiple gene regulatory proteins have been found to recognize methylated H3K4, including core components of the transcription machinery such as TAF3 (Kungulovski et al. 2016) and BPTF (Wysocka et al. 2006). As these readers have well-defined binding pockets tailored for methylated lysine, mutation of H3.2K4 would most likely lead decreased association of these reader proteins with target genes, although it is unclear why loss of these broadly acting gene regulatory factors would selectively impact Polycomb target gene expression. Interestingly, a recent study in which core components of H3K4 methyltransferase complexes were targeted for degradation proposed a role for H3K4me3 in release of paused RNA polymerase and transcriptional elongation (Wang et al. 2023). However, we find that *H3.2^K4R^* mutant cells do not exhibit a decrease in steady state H3K36me3 levels (**Figure S4**), which were globally decreased in these targeted degradation experiments (Wang et al. 2023). Our finding indicates that H3.2K4 mutation does not globally disrupt chromatin states associated with transcriptional elongation, suggesting that the phenotypes observed in the degradation studies are independent of the H3K4 methyltransferase complex’s catalytic activity. Lastly, it is also possible that H3.2K4 is not required for activation of Polycomb group target genes *per se*, but is instead required as an anti-repressor, as previously proposed for several trithorax group factors (Klymenko and Müller 2004). Here, it is important to consider that we have only disrupted H3.2K27-dependent mechanisms of Polycomb group target gene regulation in these experiments and that additional layers of target gene regulation exist, including repression through H2Aub1. Hence, an attractive model is that H3.2K4 (and/or its methylation) is necessary for counteracting repression by H2Aub1. Supporting this possibility, we find that H2Aub1 levels are unchanged in *H3.2^K4R^* cells (**Figure S2G**), suggesting that this layer of Polycomb group target gene repression remains intact. A role for H3.2K4 in counteracting H2Aub1 repression would also help to explain the specific impact of *H3.2^K4R^* mutation on Polycomb target gene expression without affecting other developmentally regulated genes (Hödl and Basler 2012).

## Materials and Methods

### *Drosophila* Stocks and Genetics

Flies were maintained at 25°C on standard corn media (Archon Scientific). Crossing schemes are depicted in **Figure S1A,D**. All histone replacement transgenes were inherited through the maternal germline unless otherwise indicated. For adult viability measurements, the number of observed progeny of the cross shown in **Table S1** were scored. Data were plotted as percentage of observed relative to Mendelian-expected ratios. For embryonic viability measurements, 100 GFP-positive progeny of the cross shown in **Table S1** were scored for hatching at 24-36 hours after egg-laying (hr AEL). The average of three biological replicates is reported. The genotypes were confirmed by PCR using primers listed in **Table S1**. Images of larvae in **Figure 1D** were captured using a Canon EOS Rebel T3i digital camera.

For mitotic clone induction, vials were heatshocked (37°C, 12 min) 24-48 hr AEL and dissected at the wandering third-instar stage or 72-96 hr AEL and dissected 48 hours after puparium formation. The *ΔHisC, FRT40A* chromosome, the *ΔHisC, UAS-YFP* chromosome, and the *ΔHisC, twi-GAL4* chromosome were gifts of Alf Herzig. The *ubiRFP, FRT40A* and *ubiGFP, FRT40A* chromosomes were obtained from the Bloomington Drosophila Stock Center (BL#34500, BL#5629). The *H3.2^WT^*, *H3.2^K27R^*, and *H3.2^K36R^* histone replacement transgenes were described previously (McKay et al. 2015). The *H3.2^K4R^*, and *H3.2^K4R,K27R^* histone replacement transgenes were generated as previously described (Meers et al. 2018) and are reported here for the first time. Transgenic lines were created at either Bestgene (Chino Hills, CA) or Genetivision (Stafford, TX) by phiC31 integration into the VK33 attP site. Array length was confirmed by either Southern blot or nanopore sequencing (Crain et al. 2024). Full genotypes of the animals analyzed are as follows:

> **Figures 1I-K, 2A, 2B, S2A, S2B, S2F, S2G, 3A, S3, 4G, 4H--**

> H3.2^WT^: *y, w hsp70-flp*; *ubiRFP, FRT40A/ΔHisC, FRT40A*; *12xH3.2^WT^/+*

> H3.2^K4R^: *y, w hsp70-flp*; *ubiRFP, FRT40A/ΔHisC, FRT40A*; *12xH3.2^K4R^/+*

> **Figure S2D, S2F --**

> H3.2^WT^: *y, w* hsp70-flp; *ubiRFP, FRT40A/ΔHisC, FRT40A*; *12xH3.2^WT^/nub^vein^-NLS-GFP*

> H3.2^K4R^: *y, w hsp70-flp*; *ubiRFP, FRT40A/ΔHisC, FRT40A*; 12xH3.2^K4R^/nub^vein^-NLS-GFP

> **Figures 3A, 4A, 4C, 4E, S4 --**

> H3.2^K27R^: *y, w hsp70-flp*; *ubiGFP, FRT40A/ΔHisC, FRT40A*; *12xH3.2^K27R^/+*

> H3.2^K27R/WT^: *y, w hsp70-flp*; *ubiGFP, FRT40A/ΔHisC, FRT40A*; *12xH3.2^K27R^/12xH3.2^WT^*

> H3.2^K36R^: *y, w hsp70-flp*; *ubiRFP, FRT40A/ΔHisC, FRT40A*; *12xH3.2^K36R^/+*

> **Figures 4A, 4C, 4E --**

> H3.2^K4R,K27R^: *y, w hsp70-flp; ubiRFP, FRT40A/ΔHisC, FRT40A*; *12xH3.2^K4R,K27R^/+*

### Cuticle Preparations

Embryos were dechorionated and mounted in Hoyer’s medium/lactic acid following standard methods. Cuticle preparations were imaged using a Leica DM5500 B microscope with Darkfield illumination under the 10x objective. 50 handpicked GFP-positive *H3.2^K4R^* embryos were analyzed for cuticle transformations.

### Peptide Synthesis

The C-terminally biotinylated H3.2 1-43 peptides were synthesized on a CEM Liberty Blue microwave peptide synthesizer, using HE-SPPS methodology (Collins et al. 2014) and Fmoc-ProTide Rink LL amide resin (loading 0.2 mmol/g). The C-terminal biotinylated lysine was introduced using Fmoc-Lys(biotin)-OH. For H3K4me3 peptide Fmoc-Lys(Me3)-OH·HCl was used to introduce methylated K4 residue. After synthesis peptide-resins were washed three times with dichloromethane, three times with methanol, and dried in a vacuum chamber overnight. The peptides were cleaved from the resin and deprotected by a 2 hr incubation with 2 mL of cleavage solution (92.5% trifluoroacetic acid, 2.5% triisopropylsilane, 2.5% ethane-1,2-dithiol, 2.5% water) and precipitated by addition into cold diethyl ether (∼30 mL). The precipitates were collected by centrifugation and washed with diethyl ether (3 x 5 mL). After the residual ether evaporated, the peptides were dissolved in water and lyophilized. The crude peptides were purified by preparative reversed-phase high performance liquid chromatography (RP HPLC) on a Waters SymmetryShield RP18 column and lyophilized. The purified peptides were characterized by matrix-assisted laser desorption/ionization time of flight mass spectrometry (MALDI TOF MS) and analytical HPLC (Millipore Chromolith RP-18e column).

### Immunofluorescence

Immunostaining of *Drosophila* wing imaginal discs, pupal wings, and embryos was performed according to standard protocols. Primary and secondary antibodies are listed in Table S5. Images were acquired using a Leica Confocal Sp8 and processed in Image J. Signal intensity measurements were made for ten clones per genotype. For Figure 4A-E, *H3.2^K27R^* (GFP-positive) and *H3.2^K4R,K27R^*(RFP-positive) tissues were mixed together prior to primary antibody addition, mounted on the same slide, and imaged using identical parameters. A region of interest square was positioned inside (GFP-negative) and outside (GFP-positive) the clone, and a ratio of signal intensity was determined after subtracting background. The normalized ratio of signal intensity was plotted relative to *H3.2^WT^* or *H3.2^K27R^* (**Table S2**). A student’s t-test was used to calculate p values.

### Western Blotting

100 GFP-positive 16-18h old 12x or 24x *H3.2^WT^* or *H3.2^K4R^* embryos were either hand-picked or collected using a Biosorter (Union Biometrica). Protein lysates were prepared as previously described (McPherson et al. 2023). Antibodies used are listed in **Table S5**. ImageQuant TL (GE Healthcare, software v8.1) densitometry analysis was used to determine total protein and band intensity. Histone PTM signal was normalized to corresponding total protein. Normalized signals from different volumes of the same lysate were averaged and set relative to the wild-type value. This process was completed for at least three biological replicates (**Table S4**).

### High throughput data

Wing imaginal disc and pupal wing FAIRE-seq z-normalized data are from GSE97956 (Uyehara et al. 2017). Wing imaginal disc H3.3 CUT&RUN data are from GSE215017 (Salzler et al. 2023).

### *In vitro* PRC2 methylation assays

Methylation assays were performed as previously described with the following modifications (Cifuentes-Rojas et al. 2014). Purified PRC2 (EED/EZH2/SUZ12/AEBP/RbAp48; *BPS Biosciences*) was titrated (0, 12.5, 25, 50, 100 nM) against H3 (1-43) peptide substrates (0.05 µg/µL of either unmodified, H3K4me3, or H3K4R) and incubated for 1 hour at 30°C following addition of 10 µM 9:1 *S*-adenosyl-L-methionine (SAM) *p*-toluenesulfonate salt (*Sigma*) to *S*-adenosyl-L-[*methyl*-^3^H]-methionine ([*methyl*-^3^H]-SAM) (*PerkinElmer*) in a reaction volume of 10 or 20 µL (in 50 mM Tris-Cl, pH 8.5, 5 mM MgCl_2_, 4 mM DTT). Following incubation, reactions were quenched with 0.25x reaction volumes of 5X SDS loading dye, separated by SDS-PAGE, and visualized by Coomassie. Histone peptide bands were excised and solubilized in 50% Solvable (*PerkinElmer*) in water for 3 hours at 50° C, transferred to scintillation vials, and 10 mL Hionic-Fluor scintillation fluid (*PerkinElmer*) was added. Samples were dark-adapted overnight, and radioactivity was quantified via liquid scintillation (*Beckman-Coulter*). Counts were normalized to background (scintillation fluid) and molar equivalents of methylation were derived from the empirically determined specific activity of the SAM. At least two independent reactions were performed (**Table S3**). A student’s t-test was used to calculate p values.

## Acknowledgements

Stocks obtained from the Bloomington Drosophila Stock Center (NIH P40OD018537) were used in this study. Antibodies obtained from the Developmental Studies Hybridoma Bank created by the NICHD of the NIH and maintained at The University of Iowa, Department of Biology (Iowa City, IA) were used in this study. FlyBase (FB2024_02) was used throughout this study. We thank Mark Peifer for assistance with interpreting embryonic cuticle patterns. We thank members of the McKay lab, Dowen lab, and UNC Histone Gene Replacement Group for experimental advice and critical feedback on the manuscript. This work was supported by NIGMS grants R35GM128851 to DJM, R35GM152103 to JMD, R35GM126900 to BDS, R35GM136435 to AGM, R35GM145258 to RJD, T32GM135128 to GCF, and R25GM055336 to CSA.

## Author contributions

CSA and DJM conceived the study. Most experiments were designed and analyzed by CSA and DJM and performed by CSA with assistance from MPL. The *in vitro* PRC2 methylation assay and peptide synthesis were designed by GCF, KK, JMD, and BDS. BRS performed the *in vitro* PRC2 methylation assay under the supervision of GCF. KK performed the peptide synthesis. CSA and DJM wrote the original draft of the manuscript. GCF, KK, MPL, JMD, BDS, AGM, and RJD reviewed and edited the manuscript. AGM, RJD, BDS, JMD, and DJM supervised the study and acquired funding.

## References

Barbour H, Daou S, Hendzel M, Affar EB. 2020. Polycomb group-mediated histone H2A monoubiquitination in epigenome regulation and nuclear processes. Nat Commun 11: 5947. doi:10.1038/s41467-020-19722-9

Beuchle D, Struhl G, Müller J. 2001. Polycomb group proteins and heritable silencing of Drosophila Hox genes. Development 128: 993–1004. doi:10.1242/dev.128.6.993

Breen TR, Harte PJ. 1993. trithorax regulates multiple homeotic genes in the bithorax and Antennapedia complexes and exerts different tissue-specific, parasegment-specific and promoter-specific effects on each. Development 117: 119–134. doi:10.1242/dev.117.1.119

Cifuentes-Rojas C, Hernandez AJ, Sarma K, Lee JT. 2014. Regulatory Interactions between RNA and Polycomb Repressive Complex 2. Molecular Cell 55: 171–185. doi:10.1016/j.molcel.2014.05.009

Coleman RT, Struhl G. 2017. Causal role for inheritance of H3K27me3 in maintaining the OFF state of a Drosophila HOX gene. Science 356: eaai8236. doi:10.1126/science.aai8236

Collins JM, Porter KA, Singh SK, Vanier GS. 2014. High-efficiency solid phase peptide synthesis (HE-SPPS). Org Lett 16: 940–943. doi:10.1021/ol4036825

Cookis T, Lydecker A, Sauer P, Kasinath V, Nogales E. 2024. Structural basis for the inhibition of PRC2 by active transcription histone posttranslational modifications. bioRxiv 2024.02.09.579730. doi:10.1101/2024.02.09.579730

Crain AT, Nevil M, Leatham-Jensen MP, Reeves KB, Matera AG, McKay DJ, Duronio RJ. 2024. Redesigning the Drosophila histone gene cluster: An improved genetic platform for spatiotemporal manipulation of histone function. bioRxiv 2024.04.25.591202. doi:10.1101/2024.04.25.591202

Dorafshan E, Kahn TG, Glotov A, Savitsky M, Schwartz YB. 2019. Genetic dissection reveals the role of Ash1 domains in counteracting polycomb repression. G3: Genes, Genomes, Genetics 9: 3801. doi:10.1534/g3.119.400579

Dorighi KM, Swigut T, Henriques T, Bhanu NV, Scruggs BS, Nady N, Still CD, Garcia BA, Adelman K, Wysocka J. 2017. Mll3 and Mll4 Facilitate Enhancer RNA Synthesis and Transcription from Promoters Independently of H3K4 Monomethylation. Molecular Cell 66: 568–576.e4. doi:10.1016/j.molcel.2017.04.018

Finogenova K, Bonnet J, Poepsel S, Schäfer IB, Finkl K, Schmid K, Litz C, Strauss M, Benda C, Müller J. 2020. Structural basis for prc2 decoding of active histone methylation marks h3k36me2/3. eLife 9: 1–30. doi:10.7554/eLife.61964.

González-Reyes A, Morata G. 1990. The developmental effect of overexpressing a Ubx product in Drosophila embryos is dependent on its interactions with other homeotic products. Cell 61: 515–522. doi:10.1016/0092-8674(90)90533-k

Günesdogan U, Jäckle H, Herzig A. 2010. A genetic system to assess in vivo the functions of histones and histone modifications in higher eukaryotes. EMBO Reports 11: 772–776. doi:10.1038/embor.2010.124

Guo Y, Flegel K, Kumar J, McKay DJ, Buttitta LA. 2016. Ecdysone signaling induces two phases of cell cycle exit in Drosophila cells. Biology Open 5: 1648–1661. doi:10.1242/bio.017525

Hödl M, Basler K. 2012. Transcription in the Absence of Histone H3.2 and H3K4 Methylation. Current Biology 22: 2253–2257. doi:10.1016/j.cub.2012.10.008

Howe FS, Fischl H, Murray SC, Mellor J. 2017. Is H3K4me3 instructive for transcription activation? Bioessays 39: 1–12. doi:10.1002/bies.201600095

Kasinath V, Beck C, Sauer P, Poepsel S, Kosmatka J, Faini M, Toso D, Aebersold R, Nogales E. 2021. JARID2 and AEBP2 regulate PRC2 in the presence of H2AK119ub1 and other histone modifications. Science 371: eabc3393. doi:10.1126/science.abc3393

Kingston RE, Tamkun JW. 2014. Transcriptional regulation by trithorax-group proteins. Cold Spring Harbor Perspectives in Biology 6: a019349. doi:10.1101/cshperspect.a019349

Klymenko T, Müller J. 2004. The histone methyltransferases Trithorax and Ash1 prevent transcriptional silencing by Polycomb group proteins. EMBO reports 5: 373–377. doi:10.1038/sj.embor.7400111

Kungulovski G, Mauser R, Reinhardt R, Jeltsch A. 2016. Application of recombinant TAF3 PHD domain instead of anti-H3K4me3 antibody. Epigenetics & Chromatin 9: 11. doi:10.1186/s13072-016-0061-9

Kuroda MI, Kang H, De S, Kassis JA. 2020. Dynamic Competition of Polycomb and Trithorax in Transcriptional Programming. Annual Review of Biochemistry 89: 235–253. doi:10.1146/annurev-biochem-120219-103641

Lifton RP, Goldberg ML, Karp RW, Hogness DS. 1978. The Organization of the Histone Genes in Drosophila melanogaster: Functional and Evolutionary Implications. Cold Spring Harb Symp Quant Biol 42: 1047–1051. doi:10.1101/sqb.1978.042.01.105

Margueron R, Justin N, Ohno K, Sharpe ML, Son J, Drury III WJ, Voigt P, Martin SR, Taylor WR, De Marco V, et al. 2009. Role of the polycomb protein EED in the propagation of repressive histone marks. Nature 461: 762–767. doi:10.1038/nature08398

Marzluff WF, Wagner EJ, Duronio RJ. 2008. Metabolism and regulation of canonical histone mRNAs: Life without a poly(A) tail. Nature Reviews Genetics 9: 843–854. doi:10.1038/nrg2438

McKay DJ, Klusza S, Penke TJR, Meers MP, Curry KP, McDaniel SL, Malek PY, Cooper SW, Tatomer DC, Lieb JD, et al. 2015. Interrogating the function of metazoan histones using engineered gene clusters. Dev Cell 32: 373–386. doi:10.1016/j.devcel.2014.12.025

McKittrick E, Gafken PR, Ahmad K, Henikoff S. 2004. Histone H3.3 is enriched in covalent modifications associated with active chromatin. Proc Natl Acad Sci U S A 101: 1525– 1530. doi:10.1073/pnas.0308092100

McPherson J-ME, Grossmann LC, Salzler HR, Armstrong RL, Kwon E, Matera AG, McKay DJ, Duronio RJ. 2023. Reduced histone gene copy number disrupts Drosophila Polycomb function. Genetics 224: iyad106. doi:10.1093/genetics/iyad106

Meers MP, Leatham-Jensen M, Penke TJR, McKay DJ, Duronio RJ, Matera AG. 2018. An Animal Model for Genetic Analysis of Multi-Gene Families: Cloning and Transgenesis of Large Tandemly Repeated Histone Gene Clusters. Methods Mol Biol 1832: 309–325. doi:10.1007/978-1-4939-8663-7_17

Morgan MAJ, Shilatifard A. 2023. Epigenetic moonlighting: Catalytic-independent functions of histone modifiers in regulating transcription. Science Advances 9: eadg6593. doi:10.1126/sciadv.adg6593

Pengelly AR, Copur Ö, Jäckle H, Herzig A, Müller J. 2013. A Histone Mutant Reproduces the Phenotype Caused by Loss of Histone-Modifying Factor Polycomb. Science 339: 698– 699. doi:10.1126/science.1231382

Poreba E, Lesniewicz K, Durzynska J. 2022. Histone-lysine N-methyltransferase 2 (KMT2) complexes - a new perspective. Mutat Res Rev Mutat Res 790: 108443. doi:10.1016/j.mrrev.2022.108443

Rickels R, Herz HM, Sze CC, Cao K, Morgan MA, Collings CK, Gause M, Takahashi YH, Wang L, Rendleman EJ, et al. 2017. Histone H3K4 monomethylation catalyzed by Trr and mammalian COMPASS-like proteins at enhancers is dispensable for development and viability. Nature Genetics 49: 1647–1653. doi:10.1038/ng.3965

Rickels R, Hu D, Collings CK, Woodfin AR, Piunti A, Mohan M, Herz HM, Kvon E, Shilatifard A. 2016. An Evolutionary Conserved Epigenetic Mark of Polycomb Response Elements Implemented by Trx/MLL/COMPASS. Molecular Cell 63: 318–328. doi:10.1016/j.molcel.2016.06.018

Salzler HR, Vandadi V, McMichael BD, Brown JC, Boerma SA, Leatham-Jensen MP, Adams KM, Meers MP, Simon JM, Duronio RJ, et al. 2023. Distinct roles for canonical and variant histone H3 lysine-36 in Polycomb silencing. Sci Adv 9: eadf2451. doi:10.1126/sciadv.adf2451

Schmitges FW, Prusty AB, Faty M, Stützer A, Lingaraju GM, Aiwazian J, Sack R, Hess D, Li L, Zhou S, et al. 2011. Histone Methylation by PRC2 Is Inhibited by Active Chromatin Marks. Molecular Cell 42: 330–341. doi:10.1016/j.molcel.2011.03.025

Tie F, Banerjee R, Saiakhova AR, Howard B, Monteith KE, Scacheri PC, Cosgrove MS, Harte PJ. 2014. Trithorax monomethylates histone H3K4 and interacts directly with CBP to promote H3K27 acetylation and antagonize polycomb silencing. Development (Cambridge) 141: 1129–1139. doi:10.1242/dev.102392

Uyehara CM, Nystrom SL, Niederhuber MJ, Leatham-Jensen M, Ma Y, Buttitta LA, McKay DJ. 2017. Hormone-dependent control of developmental timing through regulation of chromatin accessibility. Genes and Development 31: 862–875. doi:10.1101/gad.298182.117

Voigt P, LeRoy G, Drury WJ, Zee BM, Son J, Beck DB, Young NL, Garcia BA, Reinberg D. 2012. Asymmetrically Modified Nucleosomes. Cell 151: 181–193. doi:10.1016/j.cell.2012.09.002

Wang H, Fan Z, Shliaha PV, Miele M, Hendrickson RC, Jiang X, Helin K. 2023. H3K4me3 regulates RNA polymerase II promoter-proximal pause-release. Nature 615: 339–348. doi:10.1038/s41586-023-05780-8

Wysocka J, Swigut T, Xiao H, Milne TA, Kwon SY, Landry J, Kauer M, Tackett AJ, Chait BT, Badenhorst P, et al. 2006. A PHD finger of NURF couples histone H3 lysine 4 trimethylation with chromatin remodelling. Nature 442: 86–90. doi:10.1038/nature04815

Xie G, Lee J-E, Senft AD, Park Y-K, Jang Y, Chakraborty S, Thompson JJ, McKernan K, Liu C, Macfarlan TS, et al. 2023. MLL3/MLL4 methyltransferase activities control early embryonic development and embryonic stem cell differentiation in a lineage-selective manner. Nat Genet 55: 693–705. doi:10.1038/s41588-023-01356-4

Yuan W, Xu M, Huang C, Liu N, Chen S, Zhu B. 2011. H3K36 Methylation Antagonizes PRC2-mediated H3K27 Methylation *. Journal of Biological Chemistry 286: 7983–7989. doi:10.1074/jbc.M110.194027

